# Cell-type-resolved RNA-seq reveals molecular engrams of sexual experience in *Drosophila* neuromodulatory neurons

**DOI:** 10.1101/2025.10.12.681849

**Authors:** Julia Ryvkin, Anat Shmueli, Ayalla Aharony, Mali Levi, Galit Shohat-Ophir

## Abstract

Flexible behavioral responses rely on the ability of neural circuits to adapt their physiology and output to changing social and environmental contexts. Neuromodulation plays a central role in this flexibility, dynamically tuning neuronal activity and gene expression to align behavior with experience and internal states. Yet how specific experiences and motivational conditions are encoded within neuromodulatory neurons remains underexplored. Here, we show that distinct motivational outcomes drive discrete, largely non-overlapping transcriptional programs across three neuromodulatory neuronal populations in male *Drosophila* brains. Using cell-type-resolved RNA sequencing, we profiled the transcriptomes of serotonergic (Trh), octopaminergic/tyraminergic (Tdc2), and neuropeptide F receptor (NPFR) neurons under three conditions: successful mating, sexual rejection, and the absence of social or sexual interaction. Each experience induced a unique “molecular engram” within specific neuron types. NPFR neurons exhibited the strongest transcriptional remodeling following rejection, Tdc2 neurons preferentially represented the naïve-single state, and Trh neurons displayed balanced, experience-specific tuning. Shared differentially expressed genes across neuronal classes were few and often were oppositely regulated, revealing divergent circuit-specific logic. Experience triggered multilayered reprogramming across chromatin organization, RNA metabolism, translation, proteostasis and synaptic machinery, with selective recalibration of vesicle trafficking, and neuroplasticity. We further identified a compact, cross-circuit “rejection core” enriched for circadian, stress, metabolic, neuropeptidergic, and synaptic components. Together, these findings demonstrate how neuromodulatory circuits translate social experience into coordinated molecular and synaptic adaptations that enable behavioral flexibility.

## Introduction

Flexible behavioral responses, such as transitioning between mating and aggressive displays or adjusting feeding strategies according to fluctuating energy demands, are critical for animals living in environments that are constantly changing^1-4^. Such flexibility is particularly important in complex social contexts, as group-living situations are characterized by diverse and dynamic interactions that directly impact individual fitness^5-9^. Successfully navigating these environments requires animals to rapidly integrate external sensory information, such as predation threats, mating opportunities, or resource competition, with internal motivational states to execute precise behaviors that maximize survival and reproductive success^6^.

Central to this behavioral flexibility is the ability of neurons to finely regulate their physiology and output via diverse molecular programs, known as neuronal plasticity. These programs modulate critical neuronal features such as synaptic strength, intrinsic excitability, and connectivity^10-13^. A key mechanism in this respect is the regulation of gene expression programs mediated by activity-dependent transcription factors, such as CREB and MEF2, which initiate molecular cascades that sculpt the repertoire of expressed proteins, their localization, and function^14^. These transcriptional cascades not only consolidate synaptic modifications underlying learning and memory but also allow neurons to adjust their responses according to interoceptive information about the organism’s internal milieu^15^. For instance, hormonal signals, such as cortisol during stress, or metabolic regulators, such as ghrelin and leptin, trigger specific transcription factors that align neural circuit function with motivational states^6,16-22^. This integration ensures that neurons respond to external cues and internal drives, highlighting their ability to integrate multiple layers of information to fine-tune physiological and behavioral outcomes.

One of the mechanisms underlying these state-dependent transcriptional programs involves neuromodulatory circuits that relay contextual signals across distributed neural networks^23^. Neuromodulators such as dopamine, serotonin, and neuropeptide Y (NPY) dynamically adjust synaptic efficacy and network excitability, thereby shaping both immediate neural responses and long-term adaptations, as demonstrated by studies across diverse organisms ranging from nematodes to humans^12,24,25^. For instance, the fly homolog of NPY (*Drosophila NPF*) modulates several aspects of goal-directed behaviors, such as aggressive displays, investment in mating behaviors, feeding-associated actions, sleep, and sensitivity to sensory cues that predict the arrival of such goals in response to varying motivational states^26-48^. Moreover, a large body of research on flies establishes a link between transcriptional changes and adaptive behaviors, demonstrating that social interactions can shape and at the same time be shaped by dynamic gene regulation^49-61^. In this respect, we previously identified a causal link between motivational states, NPF levels, and the modulation of goal-directed behaviors in male flies. Successful mating, particularly ejaculation, induces NPF expression in male flies and reduces their motivation to consume drug rewards^33,43^. Conversely, repeated unsuccessful mating attempts due to encounters with previously mated females lower NPF levels, increase the motivation of rejected males to consume ethanol, elevate their investment in competitive behaviors, and reduce their ability to cope with stressors ^33,43^.

A common approach to identifying state-dependent changes in cellular programs is transcriptomic profiling of brain tissues. Until recent years, most transcriptomic studies have utilized whole tissues as input for bulk RNA-seq. Although this led to valuable discoveries, the approach is typically limited to identifying genes with narrow expression patterns^43,50,54,62^, whereas broadly expressed genes that are modified within a limited subset of neurons will go unnoticed, as the effect is diluted by the overall signal from other neurons. This limitation can be addressed by dividing the brain into discrete functional units and utilizing selected neuronal populations as inputs for bulk RNA-sequencing^61,63-67^. We and others have previously used this approach to map transcriptional programs across the fly brain and discovered that each neuronal population harbors a unique pattern of expressed genes^28,61,63,65,68^.

Here, we utilized neuronal population specific bulk RNAseq to profile transcriptional programs across three neuromodulatory neuron subpopulations isolated from the brains of male flies that experienced multiple events of successful mating, sexual rejection or the lack of any social/sexual interaction. Using this dataset, we characterized differentially expressed genes (DEGs) across mating experiences within each neuron population, compared transcriptional responses among neuron types to identify both common and cell-type-specific patterns, highlighted transcriptional programs activated or repressed by social/sexual experience, and identified neuromodulators and behavior-related genes whose regulation may underlie adaptive behavioral choices. We discovered discrete transcriptional signatures for rejection, successful mating, and the lack of sexual/social interaction in each neuronal population. Notably, some genes exhibited opposite patterns of regulation in different neuron types under identical conditions, suggesting that certain transcripts are regulated in a spatially and temporally precise manner, presumably serving distinct roles across neuronal circuits. This study provides a valuable resource for exploring causal links between transcriptional dynamics and adaptive behavioral responses to social challenge and stress.

## Results

### Sexual experience induces distinct transcriptional programs within neuromodulatory neurons

To explore the transcriptional programs underlying motivational states in neuromodulatory neurons, we profiled the transcriptomic signatures of brain-derived serotonergic, octopamine/tyramine neurons, and NPF-receptor neurons under three distinct sexual conditions known to modulate motivational states^9^. Uncovering the spatiotemporal regulation of gene expression requires labeling specific neuronal populations, selectively isolating their RNA, and systematically comparing gene expression profiles across these conditions. To achieve this, we employed the Isolation of Nuclei Tagged in Specific Cell Types (INTACT) method, which relies on affinity purification of genetically labeled nuclei followed by bulk RNA sequencing (RNA-seq; Fig. 1A)^28,63,69^. We expressed the nuclear tag *UNC-84-GFP* in 3 types of neuromodulatory neurons using drivers for NPFR neurons (NPFR-Gal4), serotonergic (TRH-Gal4) and octopamine/tyramine neurons (Tdc2-Gal4) (Fig.1 1B-D). Following eclosion, male flies were aged singly for four days and subsequently assigned to one of three experimental conditions: (1) “rejected,” involving interactions with non-receptive female flies for 1 hour, three times daily, over two consecutive days; (2) “mated,” involving interactions with receptive virgin females for 1 hour, three times daily, over two consecutive days; or (3) “naïve-single”, where flies remained in social isolation (Fig. 1E). At the end of the experimental phase, 100 males from each cohort were flash-frozen and processed for RNA-seq. We performed three independent biological replicates for each neuronal population and experimental condition.

**Figure. 1.**
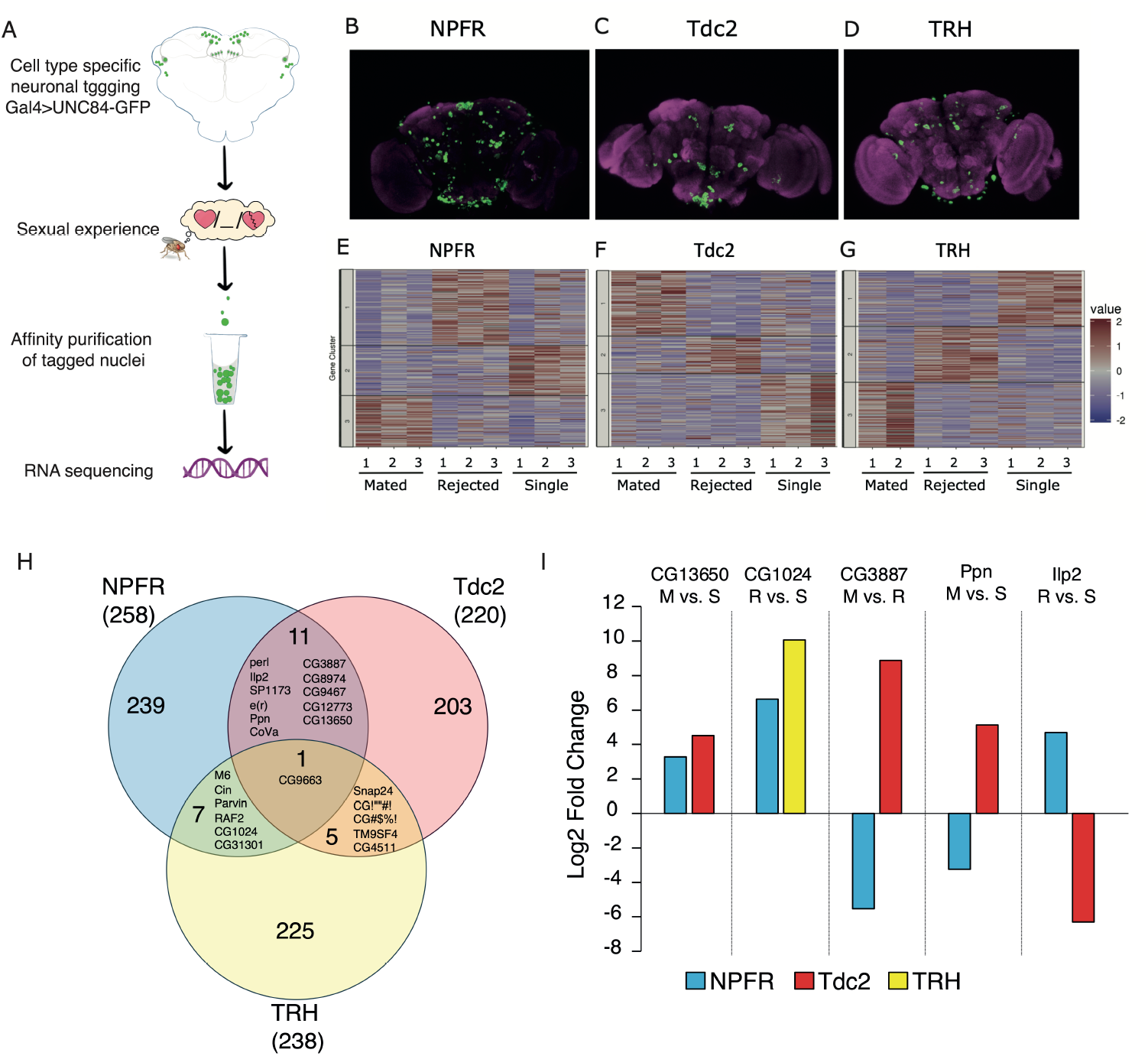
Sexual experience induces distinct transcriptional programs within neuromodulatory neurons. A. Schematic illustration of experimental pipeline. Rejected males experienced 1h long interaction with non-receptive female flies, 3 times a day for 2 days. Mated males experienced 1h long interactions with a receptive virgin female, 3 times a day for 2 days. Male flies with no sexual experience were kept in social isolation (naive-single). B-D. Expression pattern of neuronal population specific tagged nuclei using the nuclear membrane tag UNC84-GFP in NPFR (B), Tdc2 (C), and TRH neurons (D) neurons. Anti-nc-82 staining in magenta. E-G. Hierarchical clustering differentially expressed genes (DEGs) across 3 biological repreats in NPFR(E), Tdc2(F), and TRH(G) neurons isolated from brains of rejected, mated and naive-single males. Genes were clustered using Kmeans clustering method (k= 3). List of clustered genes appear in Supp table 1. H. Venn diagram of shared DEGs between NPFR, Tdc2, and TRH cell populations. I. Log2 fold change values of genes that exhibit similar and opposite regulation to the same condition across two neuronal populations.

Transcriptome analysis was conducted using pairwise comparisons among the three experimental conditions within each neuronal population, yielding three sets of differentially expressed genes (DEGs) per neuronal population (Table S1, S3). Genes significantly regulated in at least one comparison were classified as DEGs, resulting in 258 DEGs in NPFR neurons, 220 DEGs in octopamine/tyramine neurons, and 238 DEGs in serotonergic neurons (Tables S1, S3). Hierarchical clustering of these DEGs identified three major gene clusters per neuronal population, corresponding to genes preferentially upregulated in response to sexual rejection, successful mating, or lack of prior sexual or social experience (single). These clusters reveal distinct transcriptional signatures associated with each motivational state (Fig. 1E-G).

Global examination of clustering patterns revealed striking experience-dependent transcriptional organization within NPFR neurons. The rejection-associated cluster was the largest, comprising 111 DEGs, reflecting the most pronounced transcriptional response (Fig. 1E, Table S1). The naive-single cluster contained 73 genes, indicating that social isolation maintains an actively regulated transcriptional state, while the mating-associated cluster included 74 genes (Fig. 1E, Table S1). Octopamine/tyramine neurons displayed 220 DEGs, with a distribution across conditions distinct from NPFR neurons (Fig. 1F, Table S1). The largest cluster (90 genes) corresponded to genes highly expressed in naive-single males, whereas the mating-associated cluster contained 83 DEGs. In stark contrast, only 47 genes formed the rejection-associated cluster (Fig. 1F, Table S1). Serotonergic neurons exhibited 240 DEGs distributed almost evenly among the three experiences, indicative of a balanced yet distinct transcriptional response to positive sexual experience, sexual rejection, and social isolation. Specifically, mated males exhibited a cluster of 77 DEGs, while rejection yielded a similarly sized cluster of 75 DEGs (Fig. 1G, Table S1). Overall, each neuronal subtype activated a unique gene expression program across the three conditions, implying that both the valence of sexual interactions and the absence of social interactions actively drive discrete transcriptional states. Thus, the single condition represents a biologically meaningful state of social isolation rather than serving merely as a baseline.

Next, we examined whether any genes were commonly regulated across multiple neuronal populations. Analysis of shared DEGs revealed minimal overlap (only 24 shared DEGs in total), reflecting predominantly cell-type-specific transcriptional programs (Fig. 1H, I, Table S2). Remarkably, just a single gene, *CG9663*, a predicted ATPase-coupled transmembrane transporter, exhibited significant regulation across all three neuronal populations (Fig. 1H, I, Table S2). This gene exemplified cell-type specificity with striking clarity: its mRNA decreased 4.6-fold in NPFR neurons but increased 9.3-fold in serotonergic neurons following successful mating, whereas sexual rejection produced the opposite effect, inducing *CG9663* by 4.5-fold in NPFR neurons and suppressing it by 8.9-fold in octopamine/tyramine neurons (Fig. 1H, I, Table S2). Although the function of *CG9663* remains unknown, its opposing regulation reflects the precision with which individual transcripts are tailored to cell identity and motivational context.

Examining all shared DEGs further uncovered two broad regulatory patterns: consistent regulation, where genes changed in the same direction in multiple neuron types, and divergent regulation, where expression was oppositely modulated across neuronal populations (Fig. 1H, I, Supp table S2). Consistently regulated genes represented core transcriptional responses engaged by mating or rejection across neuron types. For example, *Enhancer of rudimentary* (*e(r)*)^70^ and the mitochondrial enzyme *CG9467* were consistently down-regulated in NPFR and Tdc2 neurons of mated versus rejected males (approximately −3.7 to 6.2-fold). Similarly, *CG13650* was consistently induced in NPFR and Tdc2 neurons of mated compared to naive-single males (+3.3 to +4.5-fold), signifying a shared activation of metabolic pathways in response to mating (Fig. 1H, I, Supp table S2). The gene *CG1024* (predicted to function as transcription factor) was upregulated in both NPFR and TRH neurons of rejected versus naive-single males (+6.6 to +10.1-fold), indicative of a common transcriptional response to rejection (Fig. 1H, I, Table S2). In contrast, divergently regulated genes highlighted pronounced cell-type-specific plasticity. For instance, *CG3887* (*Selenoprotein T*^*71*^) was strongly down-regulated in NPFR neurons (−5.5-fold) yet robustly up-regulated in Tdc2 neurons (+8.9-fold) in mated compared to rejected males, revealing distinct redox or stress-response programs within these neuronal populations (Fig. 1H, I, Table S2).

Similarly, the extracellular matrix protein *Papilin* (*Ppn*)^72^ decreased in NPFR neurons (−3.2-fold) but increased in Tdc2 neurons (+5.1-fold) in mated compared to single males, suggesting differential structural remodeling (Fig. 1H, I, Supp table S2). Notably, *Ilp2* (*Insulin-like peptide 2*)^73^ expression rose in NPFR neurons (+4.7-fold) but dropped substantially in Tdc2 neurons (−6.3-fold) following rejection, indicating opposing neuroendocrine tuning across circuits (Fig. 1H, I, Table S2). These findings reinforce the concept that neuronal gene expression is governed by finely tuned cell-type and experience-specific transcriptional logic rather than a global program. Moreover, they illustrate the strength of cell-type-specific RNA-seq approaches, which uncover nuanced transcriptional dynamics that would otherwise be missed (i.e., averaged out across the entire brain), obscuring significant regulation occurring in specific neuronal subsets.

### Experience-dependent remodeling of neuromodulatory neurons reveals cell-type-specific logic

Next, we examined the identified DEGs in each neuronal population. Although statistical overrepresentation analysis (GO-term analysis) did not reveal enrichment for specific biological pathways, manual analysis revealed some interesting patterns. Comparing gene expression between mated and naive-single conditions in NPFR neurons identified 111 DEGs (51 upregulated, 60 downregulated, Fig. 2A,D, Table S3), characterized by broad repression of metabolism-related transcripts such as *phosophofructokinase* (log_2_FC −3.3), and cytoskeletal and ECM genes as the *cadherin* binding protein *fat* (log_2_FC - 5.3), alongside increased expression of synaptic factors such as *Synaptotagmin*^*74*^ (log_2_FC +3.0) and Frazzled^75^ (log_2_FC +2.5). In contrast, sexual rejection elicited a distinct transcriptional response involving 74 DEGs (36 upregulated, 38 downregulated), marked by enhanced expression of couple of carbohydrate metabolism enzymes such as *Glucose dehydrogenase*^*76*^ (*Gld* log_2_FC +6.7) and mitochondrial *aconitase*^*77*^ (*Acon* log_2_FC +2.7), whereas reduced expression of proteasome related genes and oxidative stress enzymes (Fig. 2A,D). Rejection also modulated the expression of genes involved in neurite growth and synaptic remodeling, in a bidirectional manner *wengen*^*78*^(log_2_FC −2.9), *Abelson*^*79*^ (log_2_FC −5.1), *unc-5*^*80*^ (log_2_FC +5.8), and *AP-2σ*^*81*^ (log_2_FC +5.7). Additionally, rejection prominently upregulated neuroendocrine signaling factors such as *Ilp2* (log_2_FC +4.6), *Ilp5*^*82*^ (log_2_FC +6.5), and *Tachykinin*^*83*^ (log_2_FC +6.5). Direct comparison of mated versus rejected conditions yielded 122 DEGs, showing a striking bias toward downregulation (Fig. 2A, D 85 DEGs lower in mated compared to rejected, and only 37 DEGs higher). Successful mating enhanced the expression of genes involved in protein folding and synaptic assembly, including *Sorting nexin 16*^*84*^ (log_2_FC +7.0) and *Smox*^*85*^ (log_2_FC +3.3), while downregulating transcripts such as *Pinkman* (log_2_FC −4.0). The regulation of *Pinkman* is particularly intriguing, as this gene has been shown to protect neurons from synapse loss under stress conditions^86^ and to be transcriptionally induced in adult oenocytes by insulin signaling during starvation^87^. Sexual rejection maintained elevated oxidative stress responses and robustly modulated behaviorally relevant genes, including a strong downregulation of the mating-associated ubiquitin-conjugating enzyme *Ubc7* (log_2_FC −8.7) and a pronounced induction of the circadian regulator *period* (log_2_FC +7.6; Fig. 2A, D).

**Figure 2.**
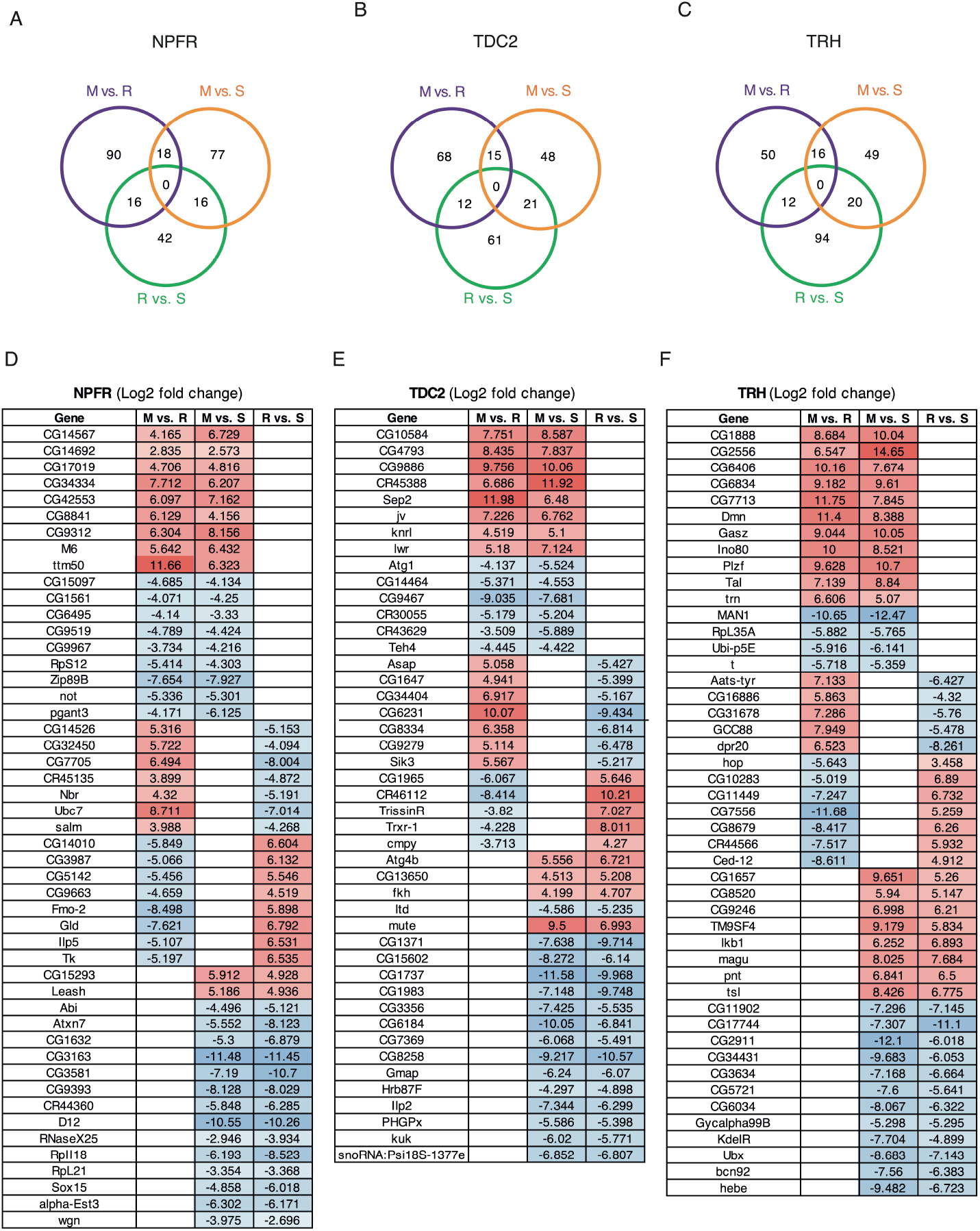
Experience-dependent remodeling of neuromodulatory neurons reveals cell-type-specific logic. A. Venn diagrams depicting differentially expressed genes that are shared across pair-wise comparisons between each two conditions within NPFR neurons (A), Tdc2 neurons (B) and TRH neurons (C). Overlapping circles indicate the number of genes which were significantly regulated in each condition. Corresponding tables summarizing Log2 fold change values of genes are significantly regulated in one social condition compared to the other two in NPFR neurons (D), Tdc2 neurons (E) and TRH neurons (F). Cell specific heat maps show the normalized fold induction of those genes

In octopamine/tyramine neurons, transcriptional responses to mating experiences differed markedly from those observed in NPFR neurons (Fig. 2B, E Table S3).

Comparing mated males to naive-single males revealed 83 DEGs (35 upregulated, 48 downregulated), characterized primarily by a broad downregulation of the protein degradation machinery and notable induction of genes related to neurotransmitter synthesis, particularly the tyramine biosynthetic enzyme *Tdc1*^*89*^ (log_2_FC +7.5). A comparison of sexually rejected males to single controls identified an even stronger bias toward downregulation (88 DEGs: 23 up, 65 down), indicating that rejection broadly suppresses gene expression in these neurons. Notably, rejection decreased the expression of genes associated with both protein folding and degradation pathways, such as *Cullin 3*^*90*^ (log_2_FC −4.1) and *Fbl6*^*91*^ (log_2_FC −4.6), while concurrently inducing oxidative stress-responsive enzymes, exemplified by *Trxr-1*^*71*^ (log_2_FC +8).

Rejection further suppressed multiple genes critical for synaptic function, including the vesicle fusion protein *Vamp7*^*92*^ (log_2_FC −11.8), the neuronal remodeling factor *orion*^*93*^ (log_2_FC −10.0), the axon guidance molecule *borderless*^*94*^ (log_2_FC −9.1), the potassium leak channel *CG1888* (log_2_FC −12.6), and the starvation-responsive kinase *Sik3*^*95*^ (log_2_FC −5.2). The shared repression of proteostasis-related genes in both mating and rejection conditions suggests that in the absence of social or sexual experience (naive-single state), octopamine/tyramine neurons maintain a higher baseline proteostatic activity. Direct comparison of mated versus rejected males yielded 94 DEGs (64 higher in mated, 30 lower), highlighting key differences attributable to rewarding versus challenging experiences. Transcripts specifically elevated in mated males included metabolic enzymes, neuronal remodeling genes (*prel*^*96*^, log_2_FC +10.7), synaptic vesicle trafficking factors (*Snap24*^*97*^, log_2_FC +6.2; *Ykt6*^*98*^, log_2_FC +7.0), and inhibitory signaling components (*GABA B R2*^*99*^, log_2_FC +4.6). Additionally, *Sik3* expression was restored following mating (log_2_FC +5.6), whereas genes elevated in rejection, such as the sleep-associated *Trissin receptor*^*100*^ (log_2_FC - 3.8) and synaptic scaffold protein *crimpy*^*101*^ (log_2_FC −3.7), were suppressed (Fig. 2B,E). Remarkably, mating also dramatically induced the molecular chaperone *Hsc70-1*^*102*^ (log_2_FC +25). Together, these results suggest that mating experience promotes a metabolically active and synaptically plastic state in octopamine/tyramine neurons, whereas sexual rejection leads to a stress-adaptive profile characterized by global transcriptional suppression.

In serotonergic (TRH) neurons, comparing mated to single males yielded 84 DEGs, with a bias toward upregulation (Fig. 2C, F Table S3, 52 genes upregulated vs. 32 downregulated). Successful mating triggered a transcriptional response characterized by increased expression of metabolic enzymes involved in lipid storage (*Ftim*^*103*^, log_2_FC +7.3), lipid catabolism (*LKB1*^*104*^, log_2_FC +6.2), and the pentose phosphate pathway (*Tal*^*105*^, log_2_FC +8.8), suggesting enhanced biosynthetic activity (Fig. 2C,F). Cytoskeletal remodeling genes were also elevated, alongside RNA regulatory factors such as *Dynactin*^*106*^ (log_2_FC +8.3) and the piRNA-associated *Gasz*^*107*^ (log_2_FC +10.0). Synaptic plasticity genes showed pronounced induction, notably *Rab3-GAP*^*108*^ (log_2_FC +23.8) and *tartan*^*109*^ (log_2_FC +5), whereas expression of *tan*^*110*^, linked to dopamine metabolism, was suppressed (log_2_FC −5.4).

Comparing rejected versus naive-single males elicited the largest transcriptional shift within TRH neurons, with 125 DEGs (62 upregulated and 63 downregulated), indicating that sexual rejection significantly impacts serotonergic neuronal transcription (Fig. 2C, F). In contrast to the metabolic activation observed after mating, rejection primarily led to reduced expression of oxidative stress genes and increased expression of components of the ubiquitin-proteasome pathway. Synaptic regulators showed mixed responses exemplified by the reduction of *HDAC6*^*112*^, *Rop*^*113*^, *twit*^*114*^, and *dpr20*^*115*^ and the upregulation the synaptic vesicle component *Snap24* (log_2_FC +4.0) and *CG9536* (log_2_FC +8.0), a protein predicted to function in retrograde vesicle transport. Notably, *Methuselah-like 10*, implicated in lifespan and starvation responses^111^, was significantly suppressed (log_2_FC −5.4). Direct comparison of mated and rejected conditions revealed 76 DEGs (48 higher in mated and 28 higher in rejected), highlighting distinct transcriptional signatures linked to these motivational states (Fig. 2C, F). Mating specifically enhanced synaptic machinery, exemplified by the induction of *dpr20* (log_2_FC +6.5), *trn*^*116*^ (log_2_FC +6.6), and *vlc*^*116*^ (log_2_FC +6.4), restored expression of the starvation-responsive kinase *Sik3*, and suppressed *cinnamon* (log_2_FC - 5.8), a gene associated with aggression^117^. Collectively, these results show that social and sexual experiences engage distinct, cell-type-specific transcriptional programs across neuromodulatory neurons. NPFR, octopaminergic, and serotonergic neurons each deploy specialized molecular responses: mating enhances metabolic, proteostatic, and synaptic remodeling pathways; rejection triggers oxidative stress, circadian, and neuroendocrine responses; and isolation maintains high baseline proteostatic and translational activity. Together, these patterns illustrate how motivational valence and social context are molecularly encoded within discrete neuromodulatory circuits.

### Systems-level, function-based analysis reveals multilayered cellular machinery underlying experience encoding

To gain a systems-level understanding of how mating, rejection, and social isolation reshape neuromodulatory circuits, we next organized each neuron’s DEGs into functional categories spanning the full spectrum of cellular machinery (Fig.3 A-C, Table S3). Rather than examining individual pairwise contrasts, this section clusters genes by their roles in mechanisms regulating gene expression spanning chromatin remodeling and transcriptional control, post-transcriptional processing (splicing, RNA localization, noncoding RNAs, piRNA), translation and tRNA charging, and post-translational modifications (phosphorylation, ubiquitination, SUMOylation), as well as core metabolic, proteostatic, and synaptic-plasticity pathways. By mapping experience-dependent transcriptional changes onto these broad regulatory layers, we reveal how discrete motivational states engage multi-level, cell-type-specific programs that coordinate gene expression, energy balance, proteome maintenance, and circuit remodeling (Fig.3 A-C). In NPFR neurons subjected to rejection, the H3K4 demethylase *Su(var)3-3*^*118*^ increased (log_2_FC +6.9), and in octopamine/tyramine neurons under rejection the MYST family histone acetyltransferase *Enok*^*119*^ rose (log_2_FC +5). In serotonergic neurons, core chromatin modifiers did not exhibit consistent shifts across conditions, but transcriptional machinery did: the basal initiation subunit *TFIIA-L*^*120*^ was elevated in mated vs. rejected TRH (+10.4), and the zinc finger regulator *PLZF*^*121*^ increased in the same comparison (log_2_FC +9.6). In NPFR neurons, selected transcription factors changed with rejection experience, including *Clock*^*122*^ (*Clk*, log_2_FC −4.1) and *Lozenge/Runx*^*123*^ (log_2_FC +6.3). Moving to RNA processing and modification, post transcriptional layers were prominently altered in TRH after mating: the *Dynactin 2, p50 subunit*^*124*^ (supports microtubule based mRNA localization) rose (log_2_FC ∼+8.3), and the piRNA pathway factor Gasz increased (log_2_FC ∼+10). Splicing factors shifted across cell types: *Slu7*^*125*^ (NPFR, mated vs. rejected) decreased (log_2_FC ∼-5.6), whereas *SF1*^*126*^ (Tdc2, rejected vs. naive-single) increased (log_2_FC ∼+6.0). Small and long noncoding RNAs were represented: *miR 4968* (NPFR, mated vs. rejected) rose (log_2_FC ∼+8.4), while the lncRNA *CR43242* decreased (log_2_FC ∼-4.8).

**Figure 3.**
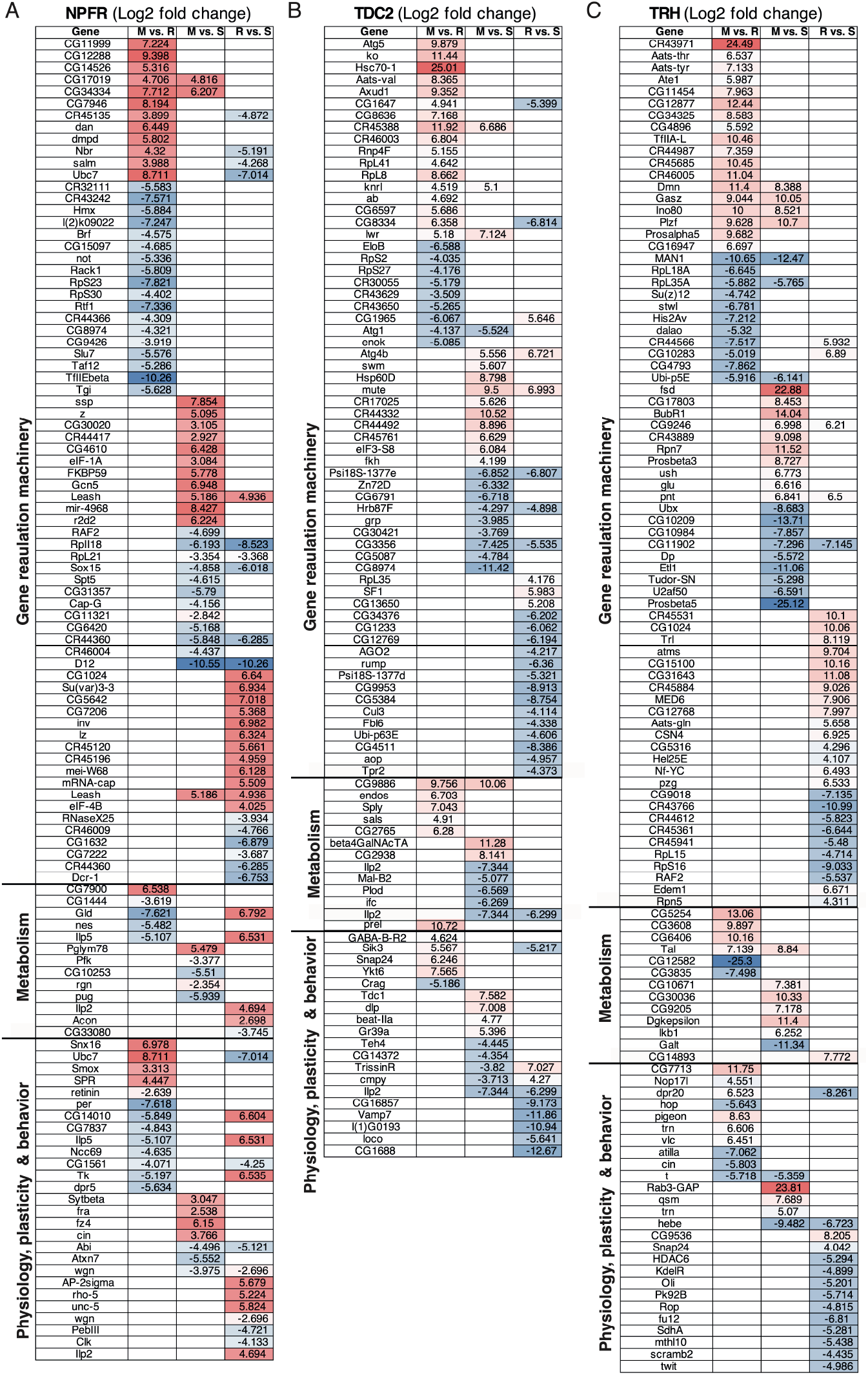
Multilayered cellular machinery encodes sexual experience. Tables depict Log2 fold change values of genes that are significantly regulated in one social condition across NPFR neurons (A), Tdc2 neurons (B) and TRH neurons (C). DEGs are divided by 3 major clusters: genes involved in cellular machineries that regulate gene expression, genes related to metabolic pathways and genes that are associated with neuronal physiology, synaptic plasticity and behavior.

As for translation regulation, enzymes for tRNA charging were induced in multiple contexts, including *Valyl tRNA synthetase* (Tdc2, mated vs. naive-single, log_2_FC ∼+8.4) and *Methionyl tRNA synthetase* (TRH, mated vs. rejected, log_2_FC ∼+10.2). Translation initiation components also shifted: *eIF3-S8*^*127*^ (Tdc2, mated vs. single) increased (log_2_FC ∼+4.9), as did *eIF-1A*^*128*^ (NPFR, mated vs. rejected, log_2_FC ∼+3.7). Ribosomal subunits showed bidirectional regulation consistent with changes in translational composition: *RpL8*^*129*^ (Tdc2, mated vs. single) rose (log_2_FC ∼+8.7), whereas *RpS16*^*130*^ (TRH, mated vs. rejected) decreased (log_2_FC ∼-9.0). In mated vs. rejected TRH, the ribosome biogenesis cofactor *Nop17l* was also higher (log_2_FC ∼+4.5). Ending with post translational modification and proteostasis, phosphorylation linked enzymes and chaperones were strongly regulated (Fig.3 A-C). The kinase *Sik3* toggled with experience in Tdc2 (rejected vs. naive-single log_2_FC ∼-5.2; mated vs. rejected log_2_FC ∼+5.6), and the major folding chaperone *Hsc70-1* was markedly induced in Tdc2, mated vs. rejected (log_2_FC ∼+25). Ubiquitin pathway components shifted in NPFR and TRH: *Ubc7* (NPFR, mated vs. rejected) decreased (log_2_FC ∼-8.7), the deubiquitinase *non-stop*^*131*^ (NPFR, mated vs. naive-single) decreased (log_2_FC ∼-5.3), the proteasome regulatory cap *Rpn7*^*132*^ (TRH, mated vs. single) increased (log_2_FC ∼+11.5), and the catalytic subunit *Prosβ5*^*133*^ (TRH, mated vs. rejected) decreased (log_2_FC ∼-25). SUMOylation capacity was represented by *lesswright/Ubc9*^*134*^ (Tdc2, mated vs. single) increasing (log_2_FC ∼+7.1), and a SUMO/ubiquitin adaptor *fate shifted*^*135*^ (TRH, mated vs. rejected) rising (log_2_FC ∼+22.9).

Sexual experience exerted a pronounced influence on genes involved in metabolism and cellular energy balance across all three neuronal subtypes (Fig.3 A-C). Carbohydrate metabolism genes were broadly upregulated following mating. For instance, the pentose phosphate pathway enzyme Transaldolase (*Tal*) was strongly induced in serotonergic neurons of mated males (log_2_FC +8.8), alongside elevated expression of the mitochondrial α-ketoglutarate transporter *CG5254* (log_2_FC +13). Octopamine/tyramine neurons and NPFR neurons from mated flies also showed increased expression of glycolytic enzymes, including glycerate kinase (*CG9886*, log_2_FC +10 in Tdc2 neurons) and phosphoglycerate mutase 1 (*Pglym78*, log_2_FC +5.5 in NPFR neurons), indicating enhanced sugar catabolism (Fig.3 A-C). Similarly, genes involved in lipid metabolism and storage were upregulated in response to mating. A monoacylglycerol lipase (*CG7900*, predicted *MAGL* ortholog) was elevated in NPFR neurons (log_2_FC +6.5), while serotonergic neurons showed increased expression of *CG10671* (a predicted lipid storage regulator, log_2_FC +7.3) and *LKB1* (a serine/threonine kinase involved in lipid catabolism and energy sensing, log_2_FC +6.2).

Genes governing neuronal physiology, synaptic transmission, and circuit remodeling underwent the most striking, experience-dependent shifts, underscoring profound plasticity following mating and rejection (Fig.3 A-C). In mated males, TRH neurons upregulated presynaptic regulators, *Rab3-GAP1* surged 23.8-fold to enhance vesicle docking and trans-synaptic signaling, while the SNARE component *Snap24* rose 4.0-fold; octopamine/tyramine neurons also boosted *Snap24* (log_2_FC +6.2) and the retrograde trafficking factor *Ykt6*^*136*^ (log_2_FC +7.0). By contrast, rejection broadly suppressed vesicle-release machinery: *Vamp7* fell 11.8-fold in Tdc2 neurons, and serotonergic neurons downregulated *Rop* (log_2_FC −4.8) and twit (log_2_FC −4.9), both essential for exocytosis (Fig.3 A-C). Experience further reshaped neurotransmitter and excitability programs: mating repressed *tan* (log_2_FC −5.4) and *cinnamon* (log_2_FC −5.8) in TRH cells, genes linked to dopamine recycling and aggression, while rejection in Tdc2 neurons silenced the potassium leak channel *CG1888* (log_2_FC −12.6), the axon-guidance molecule *borderless* (log_2_FC −9.1), and the remodeler *orion* (−10.0), collectively dampening excitability under sexual deprivation. Parallel regulation of synaptic structure reinforced these functional shifts. In Tdc2 neurons, mating induced the dendrite morphogen *perl* (log_2_FC +10.7) and the adhesion molecule *beat-IIa* (log_2_FC +4.7), while NPFR cells upregulated the membrane-trafficking protein *Sorting nexin 16* (log_2_FC +7.0) and the *SMAD-binding factor Smox* (log_2_FC +3.3). Rejection, however, activated an alternative remodeling program in NPFR neurons: the netrin receptor *unc-5* (log_2_FC +5.8) and the Dpr-interacting organizer *CG14010* (log_2_FC +6.6) were induced, even as *CG14010* and *dpr20* exhibited bidirectional tuning, upregulated by mating but repressed by rejection (Fig.3 A-C).

Finally, social isolation maintained elevated expression of developmental plasticity genes in NPFR neurons, *wengen* (+2.9) and the tyrosine kinase *Abelson* (+5.1), suggesting that the naive-single state preserves a growth-primed transcriptional landscape (Fig.3 A-C). Together, these coordinated changes in release machinery, receptor expression, and structural regulators reveal how mating and rejection sculpt distinct neuroplasticity programs across neuromodulatory circuits. Genes encoding insulin-like peptides, neuropeptide receptors, and neuromodulatory ligands showed cell type- and experience-specific expression patterns (Fig.3 A-C). In NPFR neurons, sexual rejection led to robust upregulation of *Ilp2* (+4.6) and *Ilp5* (log_2_FC +6.5), both expressed in a subset of NPFR neurons that consist of insulin-producing cells (IPCs), suggesting a stress-induced activation of insulin signaling. Conversely, in Tdc2 neurons, *Ilp2* was downregulated after mating (−6.2), indicating reduced neuroendocrine signaling in the post-mating state. The regulation of genes associated with stress response, aligns with previous findings showing that sexual rejection impairs male flies’ resilience to stressors such as starvation and oxidative challenge^33^.

Experience also affected neuropeptide receptor expression. *Trissin receptor*, a putative receptor involved in sleep and circadian modulation, was upregulated in Tdc2 neurons after rejection (+7.0) but downregulated with mating (−3.8). The neuropeptide *tachykinin* (*Tk*) was also induced in NPFR neurons following rejection (+6.5), aligning with its known role in mediating stress and social aggression. Additionally, *sex-peptide receptor* (*SPR*)^137^ was significantly upregulated in mated NPFR neurons (+4.4), consistent with its known role in regulating post-mating sleep behavior in males (Fig.3 A-C). Several experience-dependent changes pointed to broader neuroendocrine remodeling. Rejection induced *rho-5*^*138*^ (+5.2), a regulator of EGFR signaling implicated in circadian output and stress responses, and *retinin* (+2.6), a gene involved in mating behavior. Conversely, *Pinkman*, a gene linked to ecdysone signaling and male courtship, was downregulated in both mated (−4.0) and rejected (−4.7) NPFR neurons, suggesting sensitivity to both mating success and failure.

Lastly, sexual experience, particularly rejection, significantly affected genes linked to circadian regulation and sleep. In NPFR neurons, *period*^*139*^ (*per*, +7.6) and *Rnb*^*140*^ (+4.8), both core circadian regulators, were strongly upregulated in rejected males. Expression of *Trissin receptor*, modulated in Tdc2 neurons (see above), further supports the idea that mating outcome shapes circadian and sleep circuitry. Conversely, *SPR*, which promotes mating-induced sleep suppression, was elevated in mated males, suggesting opposing behavioral states driven by social success or failure (Fig.3 A-C). Taken together, these results show that mating, rejection, and isolation orchestrate coordinated, cell-type-specific gene programs across various layers, from chromatin and transcription through RNA processing, translation, proteostasis, metabolism, and synaptic machinery, thereby coupling motivational state to neuronal function. This multi-layered, neuron-resolved reprogramming provides a mechanistic bridge from social experience to circuit output and, presumably behavior.

### Courtship Failure Drives Convergent Stress‐Adaptive Programs Across NPFR, Octopamine, and Serotonin Neurons

To complement the detailed, cell type-specific pairwise comparisons and the broad, function-focused clustering of all DEGs, we next focused on genes regulated by sexual rejection across the three neuromodulatory neuron classes. Across the pairwise contrasts, rejection elicited 6 sets of DEGs, visualized in the volcano plots (Fig. 4, Table S3), that functionally converge on behaviors and physiological adaptations that are presumably relevant for coping with courtship failure.

**Figure 4.**
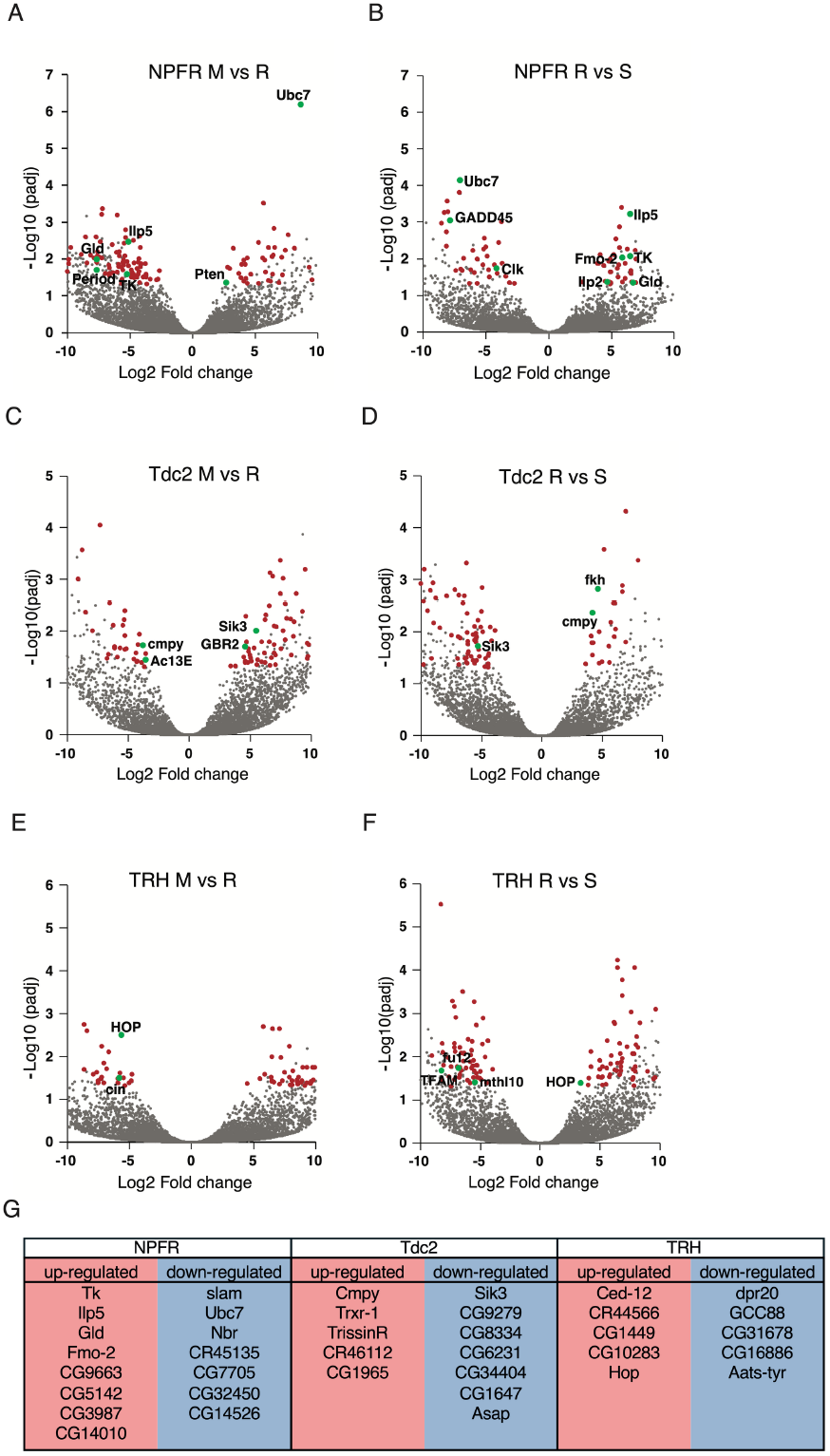
Sexual rejection induces transcriptional program within neuromodulatory neurons. Volcano plots depicting DEGs regulated by rejection in NPFR neurons (A-B), Tdc2 neuros (C-D), and TRH neurons (E-F). Grey dots represent non significantly expressed genes, red dots depict DEGs, and green dots mark DEGs of interest. G. Table summarizing rejections specific DEGs.

In NPFR neurons, the rejection-versus-mated and naive-single comparisons were dominated by induction of stress/circadian and neuroendocrine genes with concomitant downregulation of growth and proteostatic factors. Two circadian clock genes, *Clock* (*Clk*) and *period* (*per*), were differentially regulated in rejected males compared to naive-single and mated males, respectively (Fig.4A,B, Table S3). Growth arrest and DNA damage-inducible 45 (*Gadd45*)^141^, and Phosphatase and tensin homolog (*pten*)^142^ were down-regulated in rejected males compared to naive-single and mated males, respectively, both of which are related to the foxo activity pathway (Fig.4A,B, Table S3)^105,106^. In octopamine/tyramine neurons, sexual rejection was associated with transcriptional changes indicative of enhanced output. Expression of *crimpy* (*cmpy*), a dense-core vesicle protein, and Adenylyl Cyclase 13E (*AC13E*^*143*^) was increased, while the inhibitory receptor *GABA-B-R2* that is implicated in regulating lifespan, stress resistance, insulin signaling, circadian rhythms, and responses to female pheromones was downregulated (Fig. 4C,D)^69,111-113^. This combination of upregulated vesicle release machinery and cAMP signaling, together with reduced inhibitory receptor expression, suggests that sexual deprivation promotes elevated octopamine release capacity. In TRH neurons *HOPSCOTCH* (*HOP*)^144^, a tyrosine-kinase protein involved in adult circadian locomotor rhythmicity was upregulated in rejected males compared to both mated and naive-single males (Fig.4E,F)^107^. *Cinnamon* (*cin*), which is involved in inter-male aggressive behavior, was upregulated in rejected compared to mated males (Fig.4E)^108^. *1-Acylglycerol-3-phosphate O-acyltransferase 2* (*Agpat2*)^145^ and *mitochondrial transcription factor A* (*TFAM*), which are related to behavioral response to ethanol and response to oxidative stress, respectively, were downregulated in rejected compared to naive-single males (Fig.4F)^109,110^.

When lastly restricted our attention to genes that were significantly altered in rejection relative to both naive-single and mated conditions, generating a compact, rejection-specific “cores” emerged in each population that crystallize the molecular essence of sexual deprivation. In total, we found 15 genes in NPFR cells and 12 in both Tdc2 and TRH cells (Fig.4H), which were enriched or depleted in rejected males. These 3 rejection-specific modules converge functionally on four themes: genes associated with competitive behaviors and arousal states, modulation of synaptic output, and remodeling of metabolic and or stress response pathways.

The rejection-core set includes a compact block of behavior-related genes spanning aggression, arousal-circadian control, and investment in mating. In NPFR neurons, the *tachykinin* neuropeptide that is known to drive aggression and participates in the processing of anti-aphrodisiac pheromone inputs is strongly upregulated. *Ubc7*, an E2 ubiquitin-conjugating enzyme previously associated with courtship regulation^*88*^, is downregulated in NPFR neurons, pointing to potential molecular adjustments in reproductive and competitive behaviors following rejection. In the same NPFR population, *Ilp5* is induced, linking the rejection state to insulin-like signaling that can influence reproductive effort and resource allocation. Rejection also engages arousal and clock pathways represented by the upregulation of *Trissin receptor* (*TrissinR*) is up in Tdc2 neurons and of *hopscotch* in TRH neurons.

Synaptic and secretory machinery is prominently represented, indicating that these behavioral shifts are supported by adjustments in vesicle biogenesis, trafficking, and membrane organization. In Tdc2 neurons, the dense-core vesicle protein *Crimpy* is upregulated, whereas multiple trafficking and endomembrane regulators, including *Asap, CG9279, CG8334, CG6231, CG34404*, and *CG1647*^*146*^, are reduced, pointing to a coordinated retuning of release and recycling in aminergic terminals. In TRH neurons, *ced-12*^*147*^, a Rac-pathway activator linked to cytoskeletal remodeling, is increased, while the Golgi tether *GCC88*^*148*^ and the synaptic adhesion molecule *dpr20* are decreased, consistent with selective adjustments to serotonergic output and connectivity. In NPFR neurons, *slam*^*149*^ and *Nibbler*^*150*^ are reduced, suggesting altered proteostasis and RNA-processing environments that can influence neuromodulator release and signaling potency.

A third, stress and metabolism-oriented axis of the rejection core highlights the potential physiological cost of this state. Tdc2 neurons upregulate thioredoxin reductase-1, a key antioxidant enzyme, while simultaneously downregulating *Sik3*, a metabolic kinase that integrates nutrient status with cellular programs; together these changes point to redox compensation alongside altered energy signaling. NPFR neurons elevate *Fmo*-2^151^, a flavin monooxygenase implicated in xenobiotic/redox metabolism, and also increase *Gld*, consistent with adjustments in amino-acid/energy handling. In the same NPFR population, *Ilp5* is induced, linking the rejection state to shifts in insulin-like signaling that can influence both metabolism and resource allocation. In TRH neurons, *TyRS*^*152*^, a tyrosyl-tRNA synthetase, declines, hinting at selective tuning of translational capacity under rejection. Across the three neuromodulatory classes, this constellation of antioxidant, metabolic, and translational genes is consistent with the increased sensitivity to starvation and oxidative challenge characteristic of rejected males, while the behavior and synapse-related modules above outline how the same core set supports elevated aggression, altered arousal/circadian tone, and changes in mating investment.

## Discussion

In this study, we extend our previous spatial transcriptomic maps of the *Drosophila* brain, which revealed distinct transcriptional and A-to-I RNA editing patterns across neuronal populations^28,68,69^, by examining how social and sexual experiences remodel gene expression in three key neuromodulatory populations. We employed four complementary strategies: cell-type-specific pairwise comparisons, hierarchical clustering, systems-level function-based categorization, and a targeted “rejection program” analysis across all three neuron types, to reveal both highly cell-type-specific and shared stress- and reward-related transcriptional programs.

Pairwise comparisons within each cell type (mated vs. naive-single, rejected vs. naive-single, mated vs. rejected) cataloged condition-specific DEGs and provided the raw contrasts that define each motivational state. Hierarchical clustering of those DEGs within each neuronal population revealed coherent modules of genes preferentially upregulated in rejection, mating, or isolation, highlighting the relative magnitude and polarity of each cell type’s response. This global analysis demonstrated that NPFR neurons mount their strongest response to rejection, octopamine/tyramine neurons diverge most under social isolation, and serotonergic neurons respond robustly to both mating reward and rejection. Despite minimal overlap (only 24 shared DEGs). This divergent regulation of the same genes (e.g., CG3887, Papilin, Ilp2) underscores how cell identity shapes the valence‐dependent transcriptional logic.

Our systems-level, function-based analysis organized the identified DEGs into seven broad regulatory layers: chromatin modifiers, transcription factors, RNA-processing machinery, translational components, post-translational modifiers, metabolic pathways, proteostasis networks, and neuronal effectors that reshape neuronal physiology. Finally, we distilled a focused “rejection program” by selecting genes reproducibly altered in rejected males across all three neuromodulatory populations, uncovering compact, circuit-specific signatures genes regulating competitive behaviors, stress resilience and metabolic adaptation, that together define courtship failure as an active stressor. Together, these four analytical lenses provide a comprehensive picture of how sexual experience (i.e, positive, negative, or absent), is encoded into the fly brain’s transcriptome at multiple scales.

A central theme that emerges from recent studies is that aggression, courtship, feeding, and related states are not discrete, all-or-none outputs, but rather flexible and scalable behaviors shaped by neuromodulatory control. Multiple groups have identified compact circuit nodes that act as convergence points where internal state, sensory cues, and experience meet. Duistermars et al., mapped a small “threat module” where pheromonal input and internal-state signals combine to scale the intensity and persistence of aggression, rather than toggling it on/off^153^. Hoopfer et al., showed that activating P1 command neurons creates a long-lasting internal state whose expression as courtship or aggression depends on stimulus strength and context, a leaky-integrator architecture that sums inputs over time^154^. At faster timescales, Cheriyamkunnel et al., demonstrated that tyramine and octopamine, although both produced by Tdc2-labeled cells, play separable roles: tyramine rapidly gates the feeding-versus-courtship choice, while octopamine sets vigor and initiation speed^53^. Neuromodulatory control also sits at sensory entry points: Andrews et al., showed that octopamine biases Gr32a-mediated pheromonal suppression to tune aggression and courtship^155^. In parallel, serotonin sets behavioral gain in opponent fashion, as Pooryasin & Fiala identified a small serotonergic brake that imposes quiescence across locomotion, feeding, and mating, and elevating 5-HT tone can abolish the “loser effect” by recruiting defined 5-HT1B targets^156^. Finally, Zhang et al., uncovered a recurrent NPF-pCd loop that maintains mating drive over hours to days, is acutely reset at copulation by copulation-reporting neurons, and recovers under slow CREB2-dependent brakes (e.g., *Task7*), establishing a scaffold for persistent motivation^39^.

Our cell-type-specific RNA-seq maps concrete molecular levers onto this circuit logic across three neuromodulatory classes. After rejection, octopaminergic/tyraminergic neurons upregulate *crimpy* (*cmpy*), a dense-core vesicle factor, and *Adenylyl Cyclase 13E* (*AC13E*), while downregulating *GABA-B-R2*. This combination is expected to increase neuromodulator availability and cAMP signaling while relieving inhibitory tone, conditions that would lower the activation threshold and expand the dynamic range of downstream neuronal targets. In serotonergic neurons, rejection induces selective adjustments in vesicle cycling and exocytosis machinery, such as upregulation of *Snap24* and downregulation of *Vamp7* and *Rop*, suggesting a rebalancing of serotonin output that could alter how repeated social cues are processed. In addition, there are changes in circadian/kinase signaling (e.g., *Hopscotch, Cryptochrome*). NPFR neurons add a complementary layer: induction of *Tachykinin*, suppression of *Ubc7*, and changes in circadian (*per, Clk*) and metabolic (*Ilp2, Ilp5*) pathways suggest integration of arousal, time-of-day, and energy balance into competitive behaviors, presumably contributing to the NPF-pCd loop’s set-point and recovery kinetics are reset after rejection.

We previously performed metabolic analysis of male flies under the 3 conditions ^33^. Despite the dramatic behavioral differences and their sensitivity to oxidative and starvation stress, rejected males do not appear to suffer an energy deficit in baseline conditions. Instead, they exhibit more nuanced metabolic reprogramming^33^. Key indicators of energy status such as triglyceride reserves, body weight, and glucose content were indistinguishable between rejected and mated males. This means rejected flies were not energetically depleted or starving prior to the stress tests. In fact, across mated, rejected, and naive-single cohorts, there were no significant differences in total TAG levels (energy stores) or in hemolymph and whole-body glucose measurements. Thus, the heightened sensitivity of rejected males to starvation and oxidative stress cannot be explained by lower energy reserves at baseline. Comparing the metabolomic profile of male flies under the 3 conditions^33^ to the transcriptomic highlight specific metabolic pathways that are altered in rejected male flies compared to mated ones.

Notably, amino acid and heme metabolism show concordant changes at the gene and metabolite levels. Rejected males accumulate 5-aminolevulinic acid (5-ALA), a precursor in heme biosynthesis, along with elevated glycine and acetyl-glutamine, relative to mated and naive-single males. These metabolites suggest a bottleneck in heme production: if 5-ALA builds up, downstream heme synthesis may be slowed. Consistently, transcript data show upregulation of Gadd45, a stress-response gene, in rejected males (indicating cellular stress), which aligns with potential heme pathway disruption. Impaired heme synthesis could reduce heme oxygenase (HO) activity, leading to less breakdown of pro-oxidant heme byproducts. This overlap in heme/porphyrin metabolism at both metabolite (5-ALA accumulation) and gene (stress gene induction) levels supports the idea that rejected males experience a subtle oxidative stress burden, matching their greater sensitivity to oxidative challenges in vivo^33^. In line with this, *TFAM* (mitochondrial transcription factor A) was upregulated in rejected males. TFAM drives mitochondrial DNA expression and biogenesis, so its induction suggests a compensatory response to bolster mitochondrial capacity.

Another overlap is observed in lipid metabolism. While total lipid stores (triglycerides, TAG) remained unchanged between groups, a specific fatty acid stearic acid (C18:0), was significantly higher in rejected males than in mated or naive-single males. This metabolite change aligns with transcriptomic signs of altered lipid regulation. In particular, rejected males had elevated expression of *Sik3* (*Salt-Inducible Kinase 3*), a kinase known to regulate lipid and energy metabolism. The SIK3-HDAC4 signaling suppresses fat breakdown (lipolysis) during feeding and controls metabolic gene expression. The rise in stearic acid levels could indicate a shift in how lipids are mobilized or stored. In essence, both gene and metabolite data suggest that rejected males maintain a more anabolic or storage-oriented lipid metabolism profile despite their stressful social experience.

Carbohydrate metabolism and insulin signaling also show interplay between transcripts and metabolites. Rejected males upregulate Ilp2 and Ilp5, encoding insulin-like peptides, at the transcriptional level (pointing to heightened insulin pathway activity). In metabolic terms, however, blood sugar dynamics were subtly different across groups: the main circulating sugar trehalose was *lower* in isolated virgin males and higher in both mated and rejected males. Typically, high insulin signaling would drive blood sugar down by promoting uptake and storage. The observation that trehalose did not drop in rejected flies (instead, naive-singles had the lowest trehalose) suggests an alteration in insulin sensitivity or distribution. In other words, despite insulin gene upregulation, the expected metabolite outcome (lower trehalose) was not seen, an intriguing overlap that implies insulin pathway reprogramming. This could mean that rejected males have elevated insulin production (transcripts) but possibly relative insulin resistance or impaired insulin release, resulting in sustained high trehalose levels in the hemolymph. Together, these shifts maintain fuel availability for persistent behavioral drive yet limit metabolic flexibility and heightened vulnerability to starvation and oxidative stress.

Several important questions remain open following our multi‐scale analysis. First, the temporal dynamics of these transcriptional programs warrant deeper exploration. Although we profiled flies at a fixed post‐experience timepoint, it is unclear how rapidly genes such as *Rab3‐ GAP1, Ilp2*, or *Hsc70‐1* are induced following a mating or rejection event, nor how long these changes persist. Time‐course RNA-seq, sampling NPFR, Tdc2, and TRH neurons at multiple intervals after the last social interaction, could distinguish transient stress responses from more stable “molecular memories” that bias future behavior. Likewise, it will be critical to determine whether the “rejection program” can be reversed by subsequent successful mating, and conversely whether mating signatures reappear after repeated rejection, thereby mapping the plasticity and resilience of each circuit. Second, our study leaves unresolved the causal versus correlative roles of individual DEGs. While many candidates correlate with behavioral states, functional experiments are essential to test whether altering their expression in the appropriate neuron type drives changes in aggression, courtship persistence, or stress susceptibility. Spatiotemporal manipulation of the level or function of selected DEGs is necessary to assign a causal function to their regulation. Some examples illustrate both the power and the complexity of this approach. Ubc7 provides a positive case: while global Ubc7 null mutations disrupt male courtship altogether^88^, NPFR-specific knockdown paradoxically enhances social arousal and motivation^28^. This double-dissociation explains why rejection selectively reduces Ubc7 in NPFR neurons, it is an adaptive circuit-specific gain-of-function. By contrast, Tachykinin illustrates the opposite: despite its strong induction in NPFR neurons after rejection, attempts to link this change to heightened aggression or altered pheromone processing have yielded inconclusive results. These cases underscore that not all transcript changes have straightforward functional readouts, and that careful circuit-level experiments will be necessary to distinguish genuine causal levers from correlative state markers.

Third, our reliance on broad genetic drivers inevitably masks subpopulation heterogeneity. For instance, the ∼100 NPFR-expressing cells include dopaminergic, peptidergic, and fruitless-positive neurons, each potentially contributing distinct aspects of the rejection and mating programs. Single-cell or spatially resolved transcriptomic approaches, such as expansion sequencing or single-nucleus multiome profiling, will be required to map each DEG onto its precise anatomical and functional niche, thereby revealing which subcircuits govern aggression, metabolic adaptation, or circadian re-rhythming. Fourth, the integration among neuromodulatory systems remains to be elucidated. Our data show that NPFR, Tdc2, and TRH neurons each mount both unique and overlapping gene programs in response to the same social input. Yet how these circuits interact at the synaptic, paracrine, or endocrine levels to coordinate coherent behavioral outcomes is unknown. Electrophysiological recordings and circuit mapping— perhaps combining optogenetic manipulations of one cell class with calcium imaging in the others—could reveal the bidirectional influences and feedback loops that translate molecular states into system-level behavior. Finaly, although our joint RNA-seq and metabolomic results suggest tight brain-body metabolic coordination, the signaling pathways that link circuit-specific transcriptional programs to peripheral metabolism require further study. For example, does upregulation of *Ilp2/5* in NPFR neurons alter systemic insulin signaling in the fat body or gut? Does elevated *Sik3* in Tdc2 cells feedback to regulate lipid mobilization? Measuring circulating peptide levels, performing tissue-specific knockdowns, and conducting flux analyses of key metabolites (e.g., trehalose, 5-ALA, stearate) will clarify how neuromodulator circuits orchestrate whole-body metabolic reprogramming. Addressing these questions will move us from descriptive atlases toward mechanistic understanding of how social and sexual experiences are “written” into the fly brain’s transcriptome, how those molecular inscriptions govern circuit function, and ultimately how they shape adaptive behavioral trajectories.

In summary, by integrating pairwise contrasts, function-based clustering, and a focused analysis of condition-specific signatures, we provide a comprehensive blueprint of how neuromodulatory circuits transcribe sexual experience, positive, negative, or absent, discrete, multi-layered gene-expression programs. This work not only reinforces known circuit-level mechanisms of aggression, courtship, and refractoriness, but also uncovers novel molecular nodes, spanning epigenetic modifiers, RNA-processing factors, metabolic enzymes, chaperones, and synaptic regulators, through which social experience sculpts neural function and drives adaptive behavioral outcomes.

## Materials and Methods

### Sexual experience protocols

Males and females were collected within 2 h of eclosion on CO2, 3-4 days before courtship conditioning. Males were collected into narrow glass vials (VWR culture glass tubes 10X75mm) containing food and kept single housed until the conditioning. To generate mated females for the experiment, mature males were added to the females ∼16 h before the experiment. All flies were kept in the incubator at 25°C, ∼50% humidity, and light/dark of 12:12 hours. The mated females were separated from the males on the morning of the conditioning. During the conditioning, the temperature was kept at about 25°C, and humidity was ∼55%. Generation of rejected males: Individual males were placed with mated females for 3 one-h conditioning trials (separated by 1-h rests) a day for two or four consecutive days. Females were removed after each trial. Males from the rejected cohort that managed to mate and males from the mated cohort that did not end up mating during all sessions were discarded. At the end of each session, the female fly was removed, and the males that experienced rejection were kept in the original vial for one hour of rest. Males were monitored every 10 minutes to ascertain lack of mating, and when mentioned the number of males exhibiting courtship action during training sessions was documented.

### Neuronal population specific RNAseq

Cell type specific labeled nuclei were isolated using the INTACT method (Isolation of Nuclei Tagged in A specific Cell Type technique) as previously described155,156. About 100 adult male flies collected from 3-4 days F1 generation of NPFR GAL4_driver X UAS_unc84_2XGFP reporter were subjected to sexual experience for 2 days and flash frozen. Frozen heads were homogenized using 9ml of homogenization buffer (20mM β-Glycerophosphate pH7, 200mM NaCl, 2mM EDTA, 0.5% NP40 supplemented with RNAase inhibitor,10mg/ml t-RNA, 50mg/ml ultrapure BSA, 0.5mM Spermidine, 0.15mM Spermine and 140ul of carboxyl Dynabeads −270 (Invitrogen: 14305D) was added to each sample. The heads were filtered on ice by a series of mechanical grinding steps followed by filtering the homogenate using a 10um Partek filter assembly (Partek: 0400422314). After removing the carboxyl-coated Dynabeads using a magnet, the homogenate was filtered using a 1um pluriSelect filter (pluriSelect: 435000103). The liquid phase was carefully placed on a 40% optiprep cushion layer and centrifuged in a 4oC centrifuge for 30min at ∼2300Xg. The homogenate/Optiprep interface was incubated with an anti-GFP antibody (Invitrogen: G10362) and protein G Dynabeads (Invitrogen: 100-03D) for 40 minutes at 4oC. Beads were then washed once in NUN buffer (20mM β-Glycerophosphate pH7, 300mM NaCl, 1M Urea, 0.5% NP40, 2mM EDTA, 0.5mM Spermidine, 0.15mM Spermine, 1mM DTT, 1X Complete protease inhibitor, 0.075mg/ml Yeast torula RNA, 0.05Units/ml Superasin). Bead-bound nuclei were separated using a magnet stand and resuspended in 100ml of RNA extraction buffer (Picopure kit, Invitrogen # KIT0204), and RNA was extracted using the standard protocol. Cell type specific RNAseq: The NuGEN RNAseq v2 (7102-32) kit was used to prepare cDNA from the INTACT purified RNA, followed by library preparation using the SPIA - NuGEN Encore Rapid DR prep kit. Samples were sequenced on an Illumina HiSeq using single-end 60 base pair reads.

### Differential expression analysis

Reads were trimmed using cutadapt157 and mapped to Drosophila melanogaster (BDGP6) genome using STAR158 v2.4.2a (with EndToEnd option and outFilterMismatchNoverLmax was set to 0.04). Counting proceeded over genes annotated in Ensembl release 31, using htseq-count159 (intersection-strict mode). Reads overlapping exons in each gene were counted using featureCounts160, and these counts were used as input into DESeq2161. DeSeq2 function rlog(blind=FALSE) was used to calculate normalized counts with a regularized log transformation. The DESeq() and results() functions were used to calculate gene expression differences between pairs of cell types. D. Visualization of RNA-sequencing reads from the three cell types and Elav control at marker genes of the 3 groups.

## Author Contributions and Notes

G.S.O designed research, J.R, A.S, A.A and M.L performed research, J.R performed computational analysis, J.R and G.S.O analyzed data; and and J.R, A.A and G.S.O. wrote the paper.

The authors declare no conflict of interest.

This article contains supporting information online.

## Acknowledgments

We thank all members of the Shohat-Ophir lab for fruitful discussions and technical support. This work was supported by The Israel Science Foundation Grant 174/19.

---

**Supp table 1.**
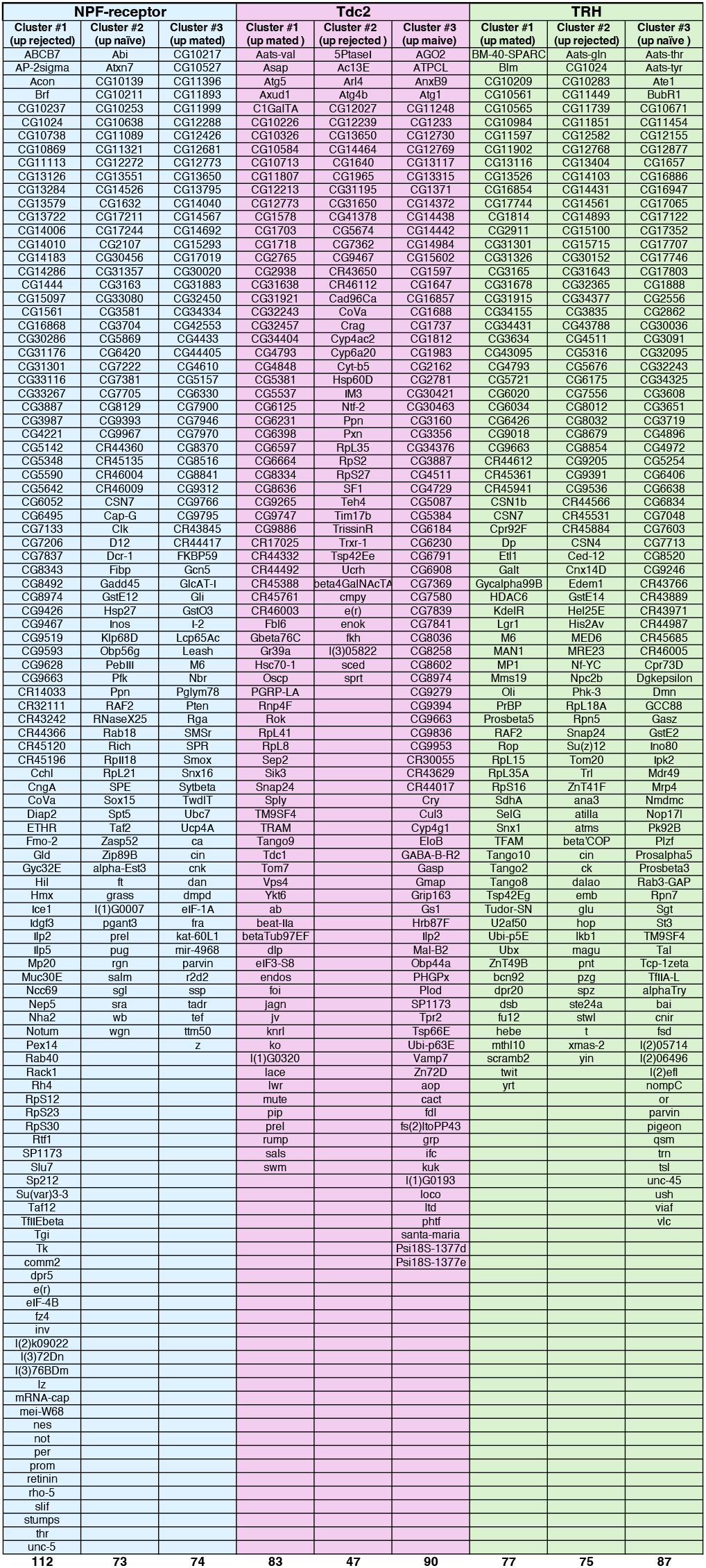
List of pairwise DEGs between two conditions across NPFR, Tdc2 and TRH neurons, that were depicted in the hierarchical clusters in Fig. 1E-F. Genes were clustered using Kmeans clustering method (k= 3).

**Supp. Table 2.** Log2 fold change values of genes that exhibit similar and opposite regulation to the same condition across two neuronal populations.

| Gene name | Log2 Fold chagne Mated vs. rejected |  |  | Log2 Fold chagne Mated vs. single |  |  | Log2 Fold chagne Rejected vs. single |  |  |
| --- | --- | --- | --- | --- | --- | --- | --- | --- | --- |
|  | NPFR | Tdc2 | TRH | NPFR | Tdc2 | TRH | NPFR | Tdc2 | TRH |
| CG3887 | -5.53 | 8.871 | 1.215 | -1.336 | 0.2404 | 3.878 | 4.194 | -8.631 | 2.663 |
| CG8974 | -4.321 | -5.936 | -0.0174 | -2.786 | -11.42 | -5.265 | 1.535 | -5.488 | -5.247 |
| CG9467 | -4.254 | -9.035 | 1.018 | -2.513 | -7.681 | 0.3591 | 1.741 | 1.354 | -0.6591 |
| CoVa | -4.846 | -2.717 | 0.9983 | -1.401 | 3.421 | 3.88 | 3.445 | 6.138 | 2.882 |
| SP1173 | -5.831 | 5.871 | 0.7744 | -2.864 | -1.936 | -0.4696 | 2.967 | -7.807 | -1.244 |
| e(r) | -3.659 | -6.182 | 1.287 | -3.243 | -4.32 | -1.694 | 0.4153 | 1.861 | -2.981 |
| CG12773 | 1.747 | 10.42 | -0.2067 | 5.933 | 1.433 | -3.5 | 4.186 | -8.983 | -3.294 |
| CG13650 | 0.06298 | -0.6952 | 0.2541 | 3.274 | 4.513 | 0.2362 | 3.211 | 5.208 | -0.01791 |
| Ppn | -0.514 | 1.514 | 1.233 | -3.231 | 5.141 | 0.5861 | -2.717 | 3.627 | -0.6464 |
| Ilp2 | -2.435 | -1.045 | -2.915 | 2.259 | -7.344 | 5.819 | 4.694 | -6.299 | 8.734 |
| prel | 6.085 | 10.72 | -2.025 | -2.2 | 3.73 | -6.451 | -8.285 | -6.987 | -4.426 |
| M6 | 5.642 | 2.491 | 3.735 | 6.432 | 2.074 | -1.271 | 0.7905 | -0.417 | -5.006 |
| CG31301 | -9.695 | -4.464 | -3.02 | -3.498 | -6.734 | -10.42 | 6.198 | -2.269 | -7.398 |
| cin | 2.949 | 2.177 | -5.803 | 3.766 | 1.874 | -2.741 | 0.8168 | -0.3025 | 3.062 |
| parvin | 4.955 | -4.292 | -0.3801 | 5.946 | 28.8 | 7.453 | 0.9902 | 33.09 | 7.833 |
| CSN7 | -8.972 | -18.47 | 1.341 | -11.82 | -16.26 | -2.62 | -2.85 | 2.21 | -3.961 |
| RAF2 | -1.794 | 1.855 | 5.159 | -4.699 | 1.702 | -0.3785 | -2.905 | -0.1532 | -5.537 |
| CG1024 | -5.326 | -2.756 | -6.319 | 1.313 | -2.619 | 3.745 | 6.64 | 0.1368 | 10.06 |
| CG32243 | 1.022 | 7.014 | 4.898 | 0.4456 | 5.391 | 6.742 | -0.5763 | -1.622 | 1.843 |
| CG4793 | 3.186 | 8.435 | -7.862 | 4.18 | 7.837 | -8.129 | 0.9941 | -0.5982 | -0.267 |
| Snap24 | 0.9779 | 6.246 | -0.5813 | -0.5995 | 2.558 | 3.46 | -1.577 | -3.688 | 4.042 |
| TM9SF4 | -6.211 | 8.642 | 3.345 | -5.967 | 3.747 | 9.179 | 0.2435 | -4.895 | 5.834 |
| CG4511 | -3.089 | 4.948 | -2.958 | 0.8606 | -3.438 | 3.48 | 3.949 | -8.386 | 6.438 |
| CG9663 | -4.659 | 3.775 | 9.394 | -0.1404 | -5.164 | 6.129 | 4.519 | -8.939 | -3.265 |

**Supp. Table 3**.

https://www.dropbox.com/scl/fi/gwz8mu7pkseo4gp7pt9qd/S3.xlsx?rlkey=ng4jrab5o2r5hmgk46cl3jt48&dl=0

